# Somatotopic organization of mechanosensory afferents in the stellate ganglion of the squid, *Euprymna*

**DOI:** 10.1101/2022.10.21.513268

**Authors:** Robyn J Crook

**Affiliations:** San Francisco State University, 1600 Holloway Avenue, San Francisco, CA 94132

## Abstract

Cephalopod molluscs are growing in popularity and use as comparative models of complex brains and behaviors. Although the gross anatomy of their central and peripheral nervous systems have been well characterized for decades, there is still very limited information about the diversity of cell types in each ganglion or lobe, their arrangement or their network properties. Unlike more standard neuroscience models, there are limited tools available for cephalopods and few validated techniques for imaging neural activity. Here, live calcium imaging in a reduced preparation of the stellate ganglion and mantle tissue reveals mechanosensory afferents and interneurons, which are arranged somatotopically in the ganglion. Retrograde labeling from stellate nerves confirms that neurons sending axonal projections to distinct dermatomes are organized in roughly oblong clusters along the dorsal side of the ganglion. This is the first demonstration of afferent somatotopy in cephalopods, and the first direct visualization of mechanoreceptive and mechano-nociceptive neurons that fire in response to localized, firm touch on the body surface. The methods and findings in this study open multiple new lines of enquiry related to sensory processing in the cephalopod nervous system.

## Introduction

Studies of the organization and function of the cephalopod nervous system have been ongoing for almost 80 years^1–9^, and its value as a comparative model of an independently evolved, complex nervous system is widely recognized^10,11^. Although there is a wealth of information from older studies about the gross structure and organization of the nervous system^12^, and a more recent explosion of molecular and genomic studies of the brain, arms and sensory structures^13–17^, functional characterization of neuronal subtypes and their network organization have remained elusive. This is in part due to the limited application to date of modern neuroscience tools that are effective in cephalopods^18^. As cephalopods continue to grow in popularity as comparative models, there is a critical need to develop improved knowledge of the fine-scale organization of the different ganglia, as well as to develop new tools to image neuronal activity in live tissue and in real time.

The cephalopod central brain is a challenging structure to study given its large size, its complexity and its largely still uncharacterized functional properties. In contrast, the smaller peripheral ganglia in the arms and mantle are attractive candidates for initial work defining neuronal subtypes based on their firing patterns. In particular, the stellate ganglion - which has received intensive but very narrowly-focused study as the location of the squid giant synapse^19–21^ – is appealing for initial characterizations of different sensory neuron sub-types given its small size, relative ease of dissection and its large nerve connections^22^. Previous work has shown that the ganglion, along with housing the large motor neurons that drive the escape jet reflex, also contains cell bodies controlling finer motor functions such as fin movement and papillae extension^23,24^, and sensory afferents responding to touch on the mantle surface^25,26^. Electrophysiological recordings from nerves carrying centripetal signal into and out of the stellate ganglion suggest that primary afferent mechanosensitive and mechano-nociceptive neuronal somata are located within the ganglion, and make their first synapses there onto interneurons that project centrally^26–28^. However, no information about their spatial arrangement, network organization or other properties has been described. In this study, a calcium imaging technique was developed to identify neurons participating in mechanosensitive pathways bringing information from the mantle, to the stellate ganglion and ultimately, to the brain.

## Methods

### Animals

*Euprymna scolopes* and *Euprymna berryi* (Fig 1A) were reared in the laboratory from captive-bred brood-stock supplied by the Marine Biological Laboratory, MA, USA. Squid were reared from eggs until early adulthood in a recirculating, artificial sweater system at 75.0 degrees Celsius. Hatchlings (0-8 weeks post-hatching) were housed in small floating enclosures within the larger, 400 US gallon tank system, and fed ad libitum on live mysid shrimp (*Amerimysis bahia*), and older juveniles and adults were fed once per day on live grass shrimp (*Paleamontes spp*.), both supplied by Aquatic Indicators, FL, USA. Squid selected for experiments were of both sexes, typically between 4-6 months post hatching, and 14-24mm mantle length. Experiments were conducted between June 2018 and March 2022.

**Figure 1:**
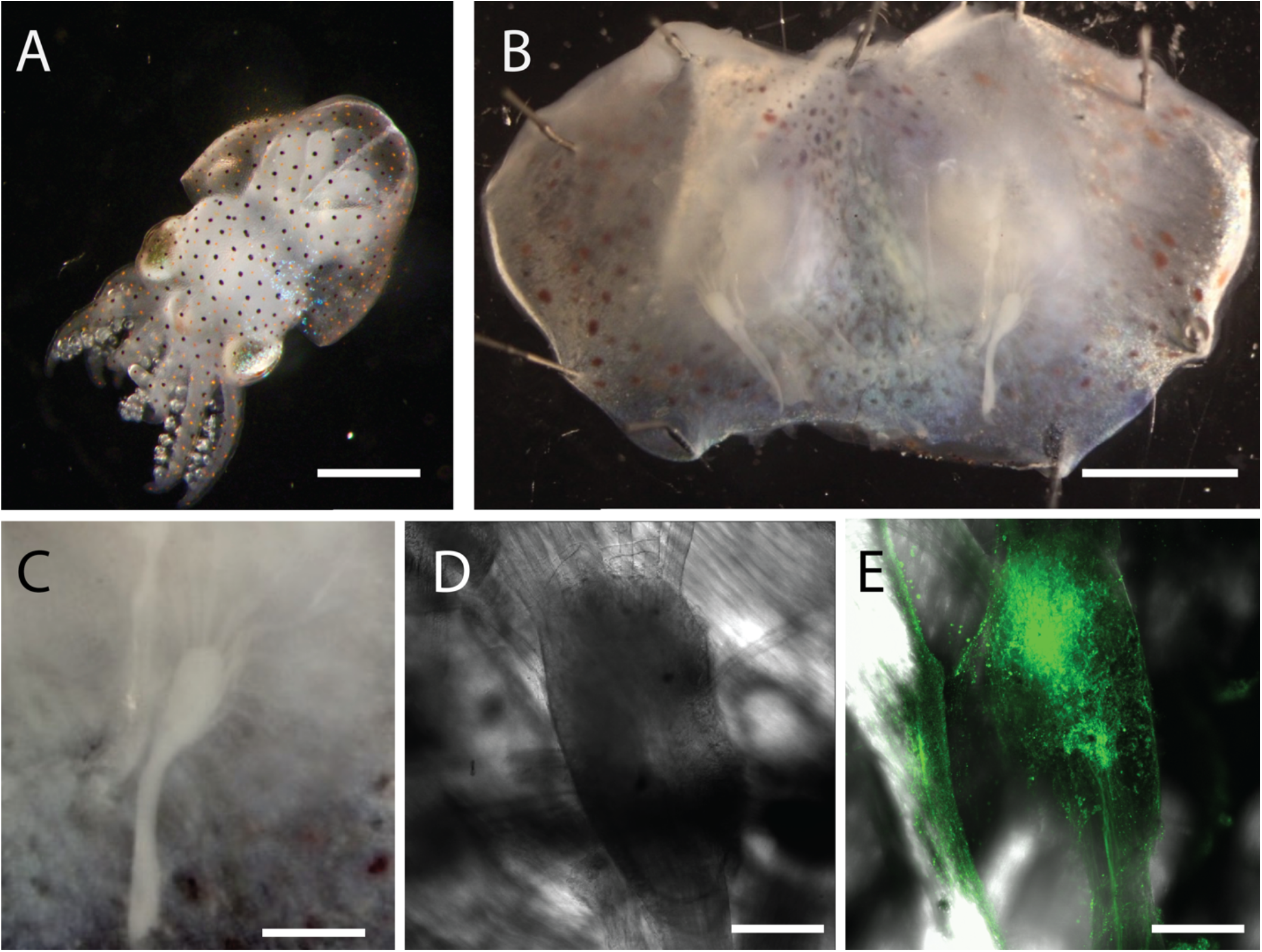
A *Euprymna berryi* juvenile. B. The mantle preparation with both stellate ganglia in their natural orientation. Imaging was conducted on the ventral surface (visible), and the dorsal surface exposed by surgically desheathing and inverting the ganglion. C. One stellate ganglion showing radiating mantle nerves, each of which innervates one mantle dermatome. D. A desheathed ganglion with individual cell bodies and nerve trunks visible E. A ganglion after injection of Cal520 dye at two locations, shown at high exposure to reveal outer extent of dye loading. Loading was heterogenous and somewhat unpredictable when dye was injected on the dorsal side.

Ethical standards and compliance: In the United States cephalopods are not included in vertebrate animal regulations that govern the use of animals in research. Although no formal approval process occurred, all animal procedures were conducted in accordance with EU Directive 63/2010/EU^29^, which contains the most stringent requirements for cephalopod research globally. Procedures, record keeping and reporting were conducted using ARRIVE guidelines^30^.

## Ca^2+^ Imaging

### Dissection

Animals were euthanized by immersion in cold, isotonic MgCl_2_ solution (330mM MgCl_2_.6H_2_0 in RODI water), until there was no evidence of reflexive movement to arm pinch and respiration had ceased for at least 5 minutes^31,32^. Squid were decapitated and decerebrated, then the mantle was pinned ventral-side-up in a Sylgard-lined dish. A ventral midline incision was made and the viscera were removed by surgical dissection. The stellate ganglion was exposed and desheathed via microdissection on the ventral (visceral cavity-facing) side, then carefully dissected away from the dorsal mantle tissue to reveal the dorsal ganglion surface. Mantle tissue was pinned tightly around the ganglion to stabilize it for imaging, and additional pins were placed at the margins of the mantle (Fig 1B). Left and right stellate ganglia (Fig 1C,D) *in situ* were loaded with dye and imaged sequentially.

### Dye preparation and loading

A stock solution of Cal520-AM dye (AAT Bioquest, catalog number 21130) was made by dissolving 50ug of powdered dye in 22.7uL of DMSO, which was stored at - 20°C. For each preparation, 5uL of stock solution was dissolved in 45uL of filtered, sterile, artificial seawater containing 0.12% pluronic acid (Pluronic F-127, Sigma, cat no. P2443), modified from^33^. 1-2uL of working concentration dye solution was injected into the cell body layer of the stellate ganglion in multiple locations for each experiment (see Fig 1E), using a glass capillary tube attached to syringe pump injection system. Dye was injected for 5-10 minutes per location and then allowed to load into the cells over the course of 1-1.5 hours after injection, during which time the dish was perfused with ASW and kept in the dark. Imaging of each ganglion occurred between 2-3 hours after dye injection. Electrophysiological recordings made from the cut end of the pallial nerve in dye-loaded preps confirmed that the tissue was healthy and there was strong evoked mechanosensory activity in response to touch on the mantle margin.

### Imaging

Ganglia were imaged at 10x on a Zeiss 710 confocal microscope. Because the ganglion surface is strongly curved, for each timepoint a z-stack of images was acquired. This greatly limited the temporal resolution of the recorded data, but ensured that activity in all cells across the surface of the ganglion was captured after each stimulation.

Each experiment followed the same procedure. A position on the ganglion was chosen for imaging, and a time series of 4-6 z-stacked images was taken at 20, 45 second intervals. The first 10 timepoints contained no mechanical stimulation of the tissue. The second series of 10 images, collected immediately after the first ten, included touches with a calibrated probe on different positions of the mantle, at timepoints 12, 14, 16, 18 and 20. In total, 42 ganglia were imaged, typically in 3-4 locations per ganglion. Of these, 27 were imaged from the ventral side of the ganglion and 15 from the dorsal side.

### Analysis

Image sequences were processed in FIJI and Matlab. First each file was cropped, adjusted for overall brightness and contrast, and reviewed manually for evidence of either spontaneous or evoked firing of neurons in the imaged region. For preparations with evidence of activity (32 of the 42 preparations), a maximum intensity projection was made from the slices, and the file was exported at a tiff sequence to Matlab. The *EZCalcium* toolbox was used to identify cells firing in response to touch on the body surface at different positions. Files were motion corrected using the default settings and ROI detection was conducted without the Neuron Classifier option, with manual curation and an estimated ROI diameter of 6-8 pixels, depending on the preparation. First, heatmaps were plotted to identify image sequences showing evidence for greater fluorescence transients in the second half of the recording, suggesting activity was enhanced by touch (Figure 2). In imaging sequences showing hotter colors in the second half of the recording, individual traces were visualized in *EZCalcium* and each identified cell was then classified into a putative primary afferent mechanoreceptor (a cell that fired in response to touch only on one bodily region), a putative mechanosensory interneuron (a cell that fired in response to touch on two or more bodily regions not likely innervated by the same primary afferent neuron), and cells that fired in patterns not associated with touch. Example traces are shown in Figure 3. Only excitatory neurons (increase in fluorescence intensity) were classified, and no attempt was made to identify possible inhibitory neurons.

**Figure 2.**
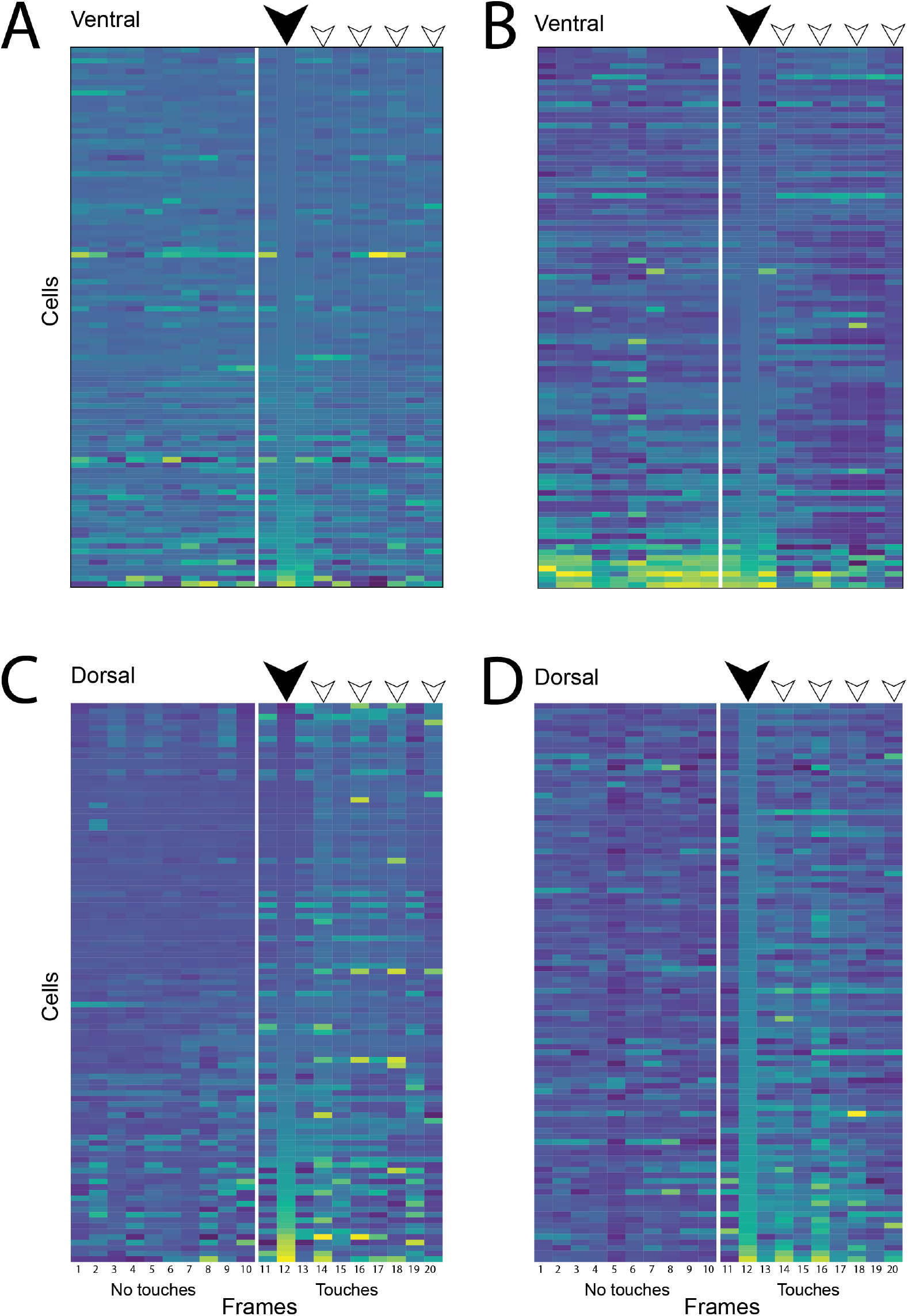
Heatmaps of normalized fluorescence values for each ROI and each imaging frame, as extracted by *EZCalcium* for Matlab. Heatmaps are shown for the 100 cells (y axis) that showed the highest brightness values in response to touch at the first stimulation location (denoted by the large, filled arrowhead at timepoint 12 on the x-axis. Further stimulations are denoted by open arrowheads at timepoints 14, 16, 18 and 20. A and B show heatmaps from imaging locations on the ventral (innermost) ganglion surface. C&D show two different examples from imaging on the dorsal surface of the ganglion. A and B show multiple cells firing during the imaging sequence but no evidence of hotter values on the right side, corresponding to touches made in sequence on the mantle surface during imaging. C and D shoe considerably hotter values on the right side, indicating that this imaging sequence contains cells responding to touch.

**Figure 3.**
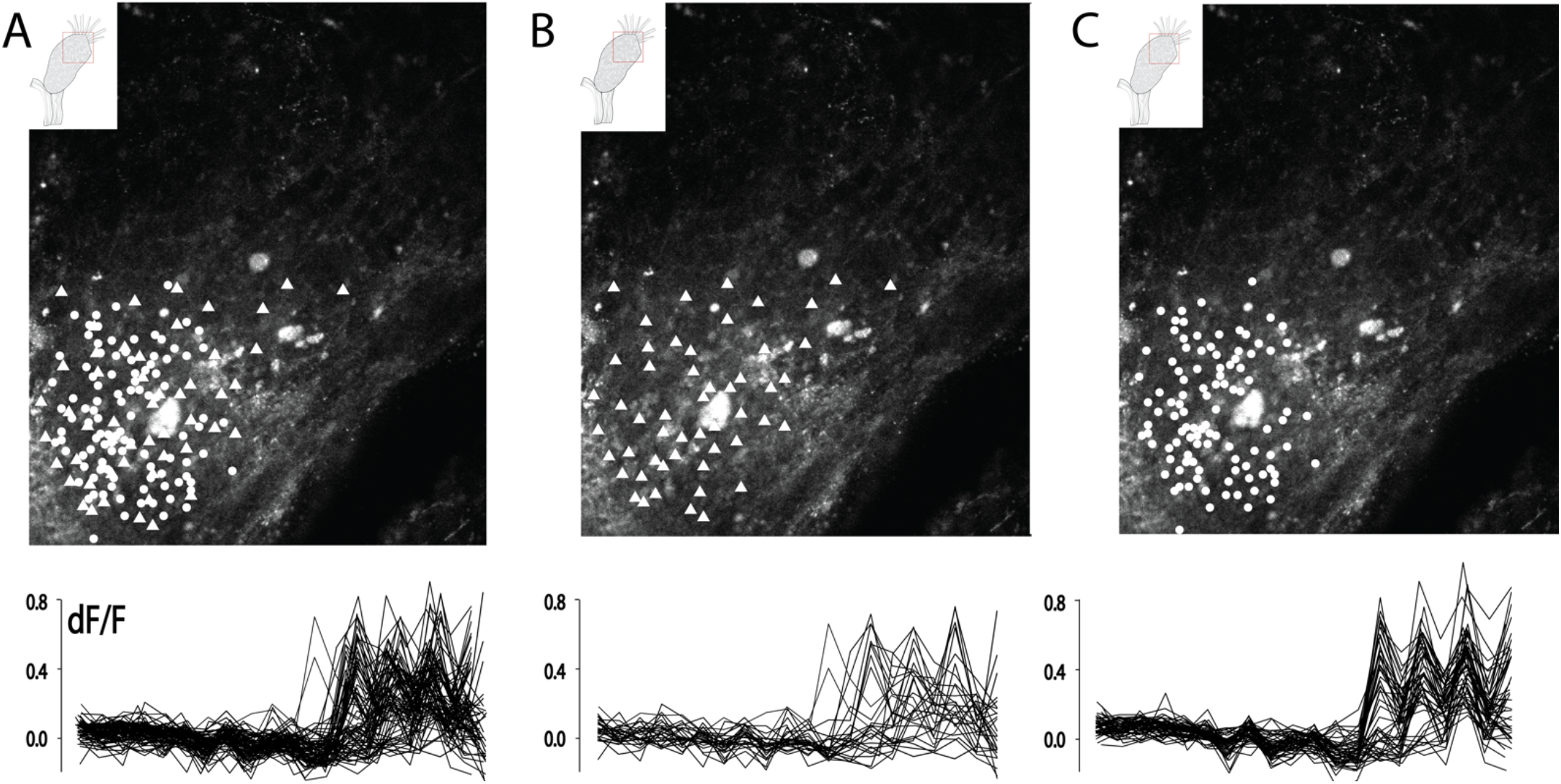
A. The dorsal-caudal region of the SG, with all cells firing in response to touch shown in white. Fluorescence transients for all cells active in response to touch shown below. B. Cells firing only in response to touch on a single location on the mantle are distributed across the ganglion, and are relatively few in number. C. Cells firing in response to touch on multiple locations are more numerous and are generally intermingled with those responding only at a single touch location.

## Neurobiotin labeling of mantle nerves

To identify the locations of neuronal cell bodies with axons projecting to the mantle muscle and skin, selected stellate nerves were backfilled with neurobiotin, following the procedure in^34^. Briefly, squid were euthanized and dissected as described above, then the stellate ganglion was surgically dissected away from the underlying muscle along with 3-4mm of the stellar nerves. One nerve from each ganglion was selected for labeling. Ganglia were washed in fASW, then the cut end of one nerve was drawn into a silicon grease well in the center of a small petri dish. The ganglion and unlabeled nerves were immersed in fASW on the outside of the well. The cut end was first immersed in RODI water for 2 minutes then the water was replaced by 5% NB in RODI water.. Ganglia were left to incubate overnight at 4degC, before the neurobiotin solution was washed off and ganglia were fixed for 2-3 hours in 4% PFA in SW on a rocking shaker, and stored for 1-2 weeks at 4C. Ganglia were then washed, permeabilized and labeled according to, then cleared for imaging following the procedure in^35^. Ganglia were imaged at 40x on a Zeiss 710 LSM confocal and image files were further processed in FIJI to adjust overall contrast and brightness.

## Results

### General procedural outcomes

Injected calcium dye successfully permeated neuronal cell bodies in the ganglia within 30 minutes to one hour after injection into the central neuropil layer, although the distribution of dye and its intensity varied considerably among different preparations (see Figure 1E for example). No obvious improvements in dye loading or spread of the dye beyond the injection location were observed in preparations allowed to load for longer than 2 hours and up to 4 hours. In all preparations, afferent signal was recorded briefly using a suction electrode on the cut end of the pallial nerve prior to imaging, and six of 42 preparations showed clearly impaired signal and were discarded. However, in some cases where strong electrophysiological signal was apparent and the imaging location was correct for capturing mechanoafferent activity, no cells showed changes in fluorescence during the imaging phase, either as spontaneous activity or activity in response to touch on the mantle surface. Given the superficial location of the mechanosensitive afferents and the need to surgically desheath the ganglion to permit imaging, it was likely these cells were damaged during dissections.

Stellate ganglia were imaged in multiple locations over the course of the study. Imaging success overall was low, with only 10 dorsal-side and 14 ventral-side locations yielding good evidence of activity during imaging, which was defined arbitrarily as at least 10 individual neurons in the field of view showing clear changes in fluorescence intensity either spontaneously or in response to touch. Post-imaging analysis suggested that the primary cause of poor neuronal activity was variations in de-sheathing success on the dorsal ganglion surface; the ganglion is tightly sheathed and connected to the inner wall of the mantle, and the mechanoreceptive neurons appear to be located right on the surface of the ganglion. Thus they are particularly vulnerable to damage during dissection and de-sheathing. Ongoing efforts in our laboratory to refine the dissection procedure may improve the success rate in future.

### Identification of evoked mechanosensory activity

Imaging sequences were assessed in real time by the experimenter for evidence of evoked responses. These were found reliably in the distal end of the dorsal surface of the ganglion (see videos S1 and S2). Although it is possible that disorganized and sparse evoked firing occurred in other ganglion regions (see videos S3 and S4), here the focus was on defining clustered organization of sensory afferents, and an exhaustive survey of all neurons participating in the mechanosensory pathway was not attempted. Fluorescence intensities from imaging sequences were extracted and plotted as heat maps where each imaging frame is plotted along the x-axis (Figure 2), organized by intensity after the first touch on the mantle with the probe (large back arrowhead). Heat maps show clearly that “hotter” colors are located on the rightmost half of the plots from imaging sequences taken from the dorsal/distal ganglion region, whereas those made from sequences taken on various locations on the ventral surface show a number of active cells but no clear difference in intensity on either side of the plot. Thus the majority of clustered mechanosensory neurons appear to be located on the dorsal/distal ganglion surface.

### Categorization of primary afferents and second-order interneurons

Neurons were clustered according to their response properties and plotted onto still frames from their imaging session to reveal spatial arrangements (Fig 3A and Fig S1). Cells were defined as probable primary mechano-afferents if their response was specific to touch on a single location on the mantle (Figure 3B), or as probably mechanosensory interneurons, if they responded to touch on multiple locations (Figure 3C).

For putative primary afferents, cells were color coded according to the location of touch on the body surface on the flat-mantle preparation that was used for imaging, which corresponded to distinct dermatomes innervated by different mantle nerves emanating from the stellate ganglion(Fig 4A&B). These dermatomes were overlaid onto the mantle of an intact squid to show their natural arrangement (Figure 4C). Examples of fluorescence traces for a subset of plotted cells are shown in 4D, separated and color coded according to the touch location where they responded. The number of primary afferents responding to touch in a given location varied among preparations, from 2-3 up to a maximum of 17. Stimulations were stippled over 6-8 mm of tissue 12 seconds prior to an image being taken, in an attempt to maximize the number of cells firing, suggesting an innervation density of 2-3 neurons per sq mm of tissue. Finer-scale mapping was not attempted in this study.

**Figure 4.**
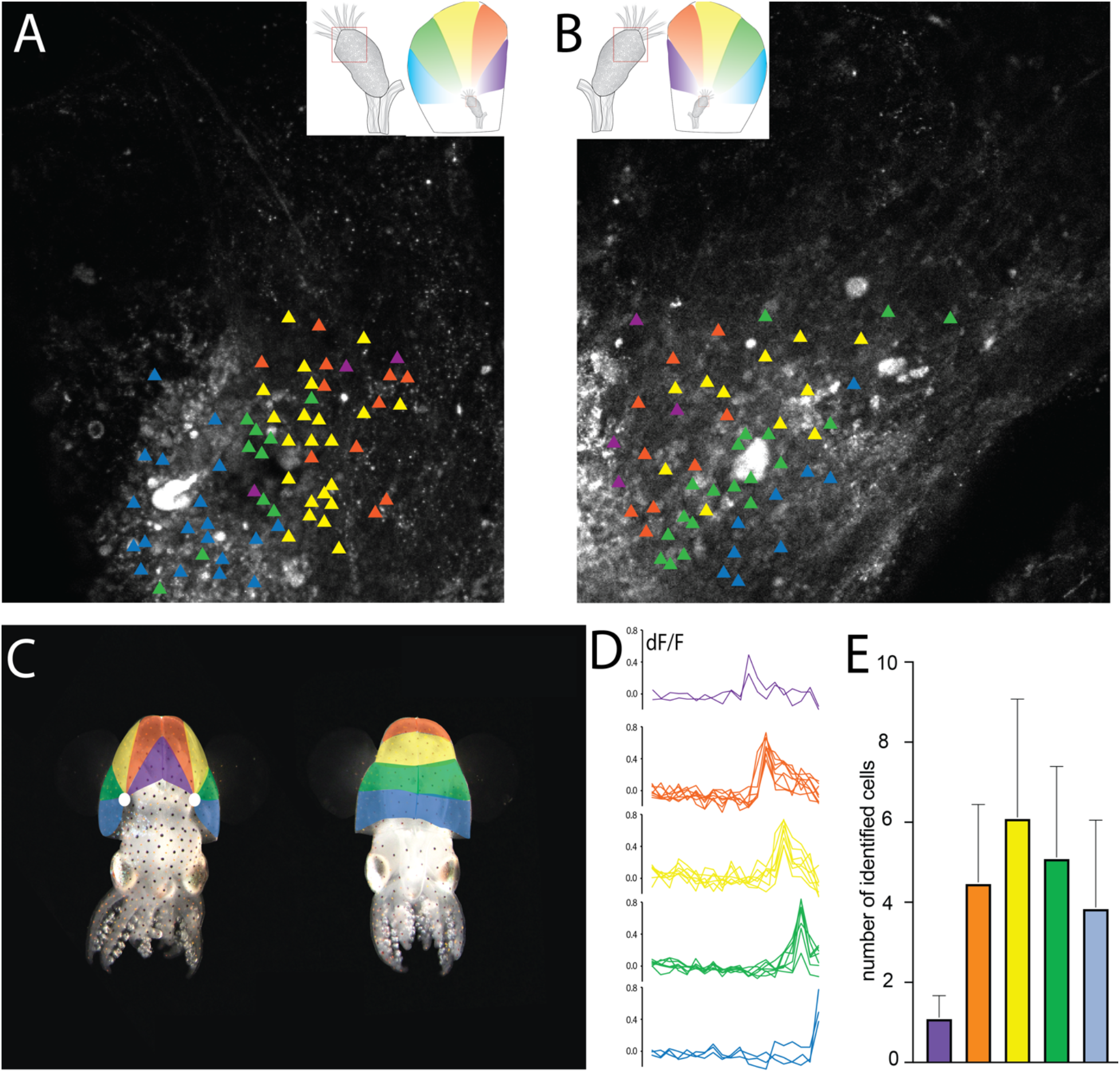
Mapping putative primary afferent mechanosensory neurons in the stellate ganglion. A and B show two different ganglion preparations, corresponding to Supplementary video files S1 and S2. Putative primary afferents are mapped as in Figure 2, but are color coded according to the location of the touch that directly preceded the imaging frame. Insets of A and B show the location on the flat-mantle preparation as pinned during imaging and stimulations. C. Dermatomes shown inset in A and B, mapped onto the squid body. Left image shows the dorsal side and right shows the ventral side. Position of the SG is denoted by a white circle, and dermatomes radiate from the ganglion in elongated wedges, converging at the ventral midline. The dorsal-rostral aspect of the mantle was not mapped due to nerves being cut here to permit inversion of the ganglion for imaging on the dorsal surface. D. Example traces of fluorescence transients for cells mapped in A and B. E. Average numbers of neurons with receptive fields in each of the colored dermatomes, taken from a total of 14 separate preparations.

For putative interneurons (cells firing in response to touch at multiple distinct locations), mapping and color coding was made more complex by the wider range of possible response patterns. Groupings were based on a combination of how broad the responses was of a given neuron (response to touch from two up to all five locations) and on the spatial character of the cell’s response. Overlays on the same still frames as in Figure 4A and B are shown in Figure 5A and 5B. Somatotopy was less clear overall, although there are some patterns evident. Most of the cells responded to touch on 3 or 4 different locations. Some cells showed responses to either the most lateral or the most central positions, suggesting that there is organization within this population of cells although it is less strict that for primary mechano-afferents. Plots of the response broadness (Figure S2) show no clear organization. Overall the impression is of loose somatotopy but less clear than for putative primary afferents

**Figure 5.**
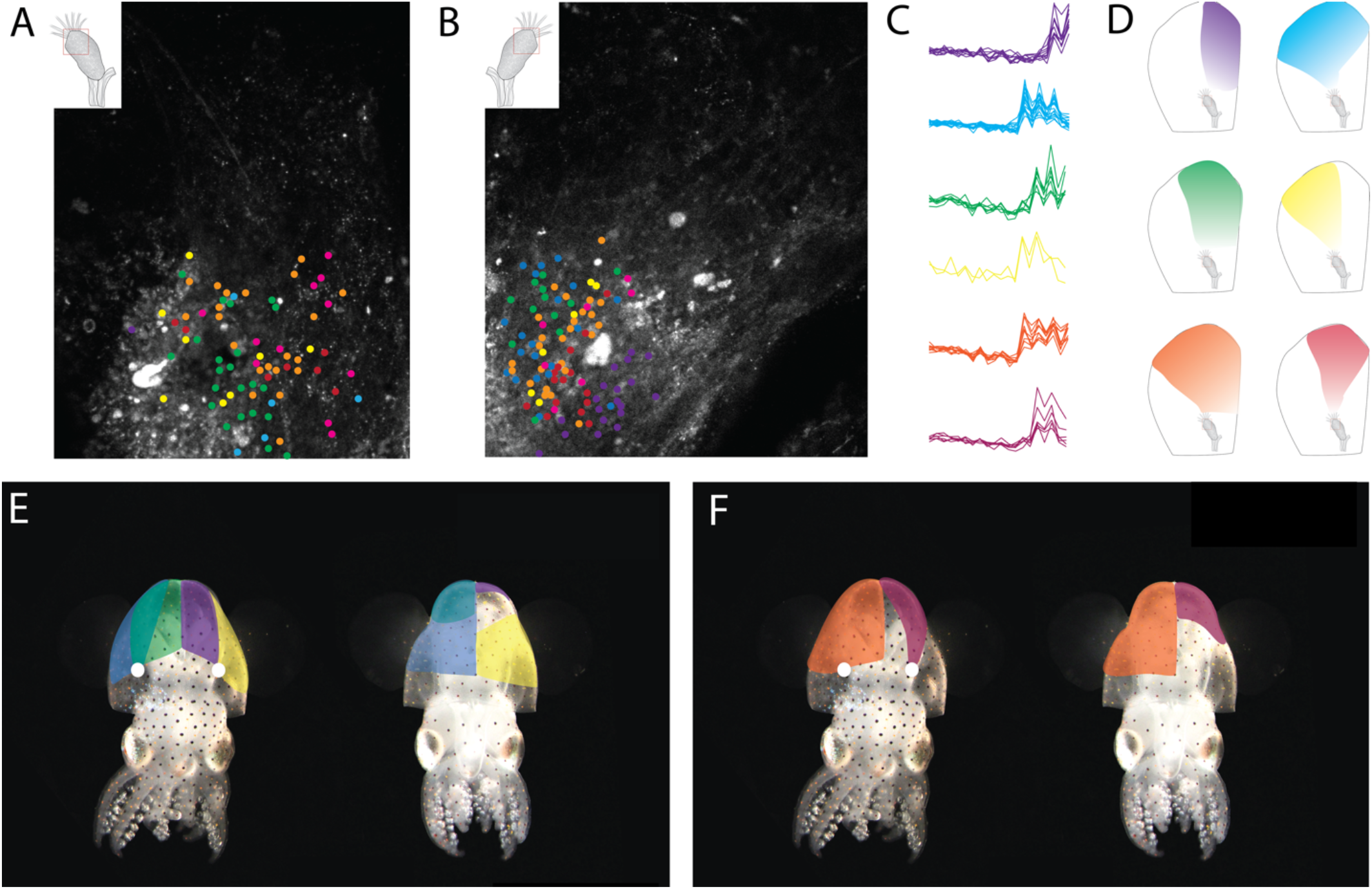
Mapping interneuron-type cells in the stellate ganglion. A and B show sill frames from the same preparation as in Figures 3&4, with cells colored according to their spatial response patterns. C. Extracted fluorescence traces for each labeled cell are shown grouped anc colored by their activation profile. D. The spatial response patterns of each group of colored cells. E. and F show the spatial arrangement of each class of cells (grouped by color), but mapped onto the mantle in a live squid rather than the flat-mantle preparation in D. A dorsal (left) and ventral (right) view of the squid is shown. Due to the complex overlap of the different spatial arrangements, the six classes of cells are shown across two panels.

### Retrograde labeling of somata with axons projecting to distinct dermatomes

Labeling generally confirmed the spatial organization of cells with axons projecting into stellate nerves around the ganglion, with lateral nerves having cell bodies along the edges of the ganglion’s dorsal surface and more central nerves having cells projecting into the midline of the ganglion (Figure 6). A number of very fine nerves emanating from the dorsal surface of the ganglion or close to the rostral end (closest to the mantle collar) were not successfully filled, and the location of cells projecting axons into the smaller nerves is not clear. These data generally support the findings from the calcium imaging experiments that primary afferent neurons tend to be located in longitudinal columns extending from the base of the stellar nerve down to the mid-point of the gangion’s rostral-caudal axis, and arranged across the ganglion surface matching the position of the emerging stellar nerve. Sizes of the filled neuronal somata and the axon diameters were variable, indicating - as expected - that the nerves contain axons belonging to a mixed population of neuronal subtypes, only a small number of which are mechanosensory afferents.

**Figure 6.**
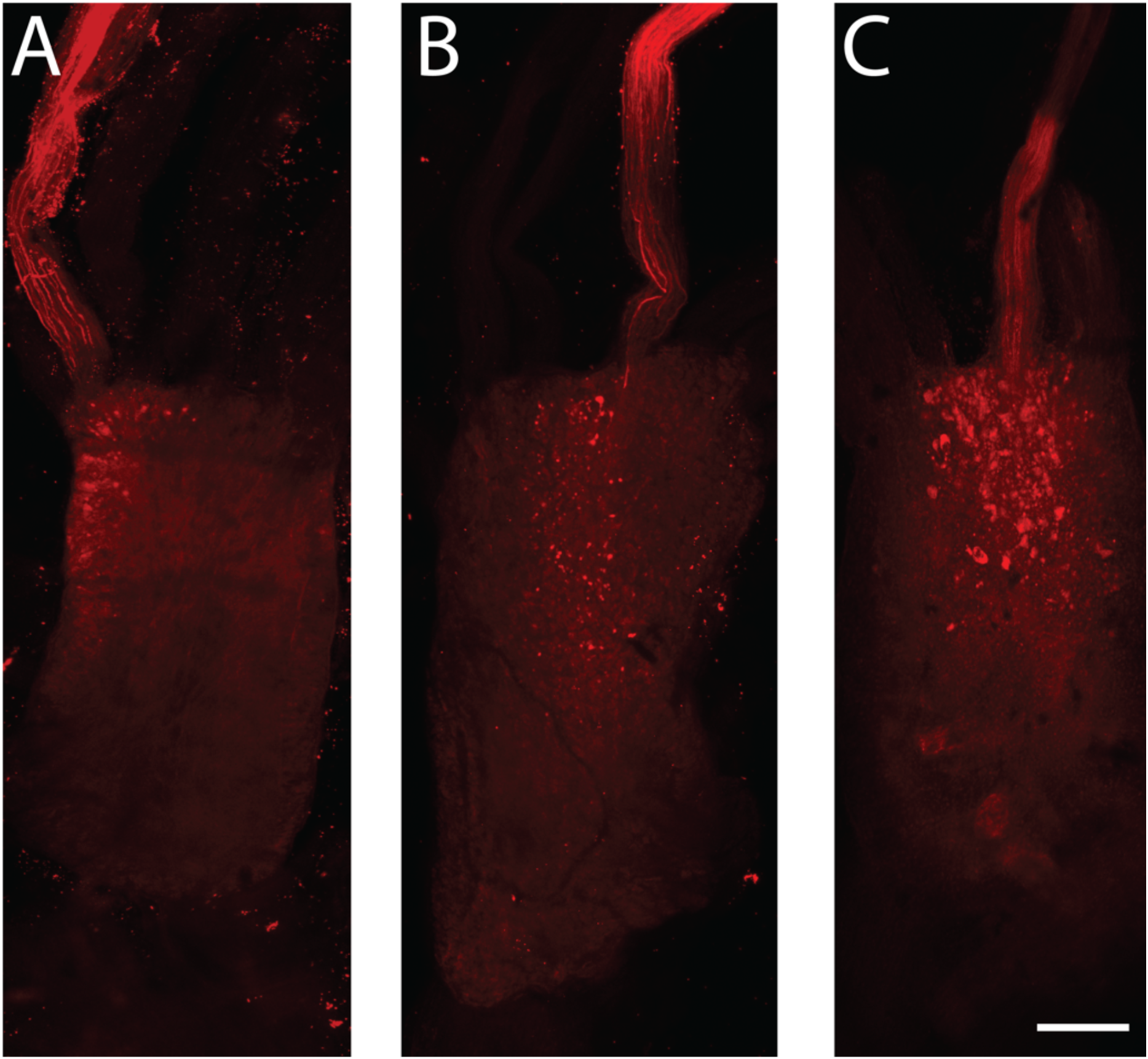
Examples of neurobiotin backfilling from three distinct mantle nerves. A. a lateral nerve projecting around the mantle and toward the ventral midline, B. a nerve projecting to the caudal mantle tip. C. a nerve projecting to the medial (midline) dorsal mantle.

## Discussion

This study provides the first functional evidence for somatotopy in the cephalopod nervous system, and reveals for the first time the peripheral ganglionic location of two related classes of sensory neuron – mechanoreceptors and mechano-nociceptors. Although fine-scale relationships between putative primary afferent neurons and their higher-order network counterparts were not clear from this initial study, it is likely that the stellate ganglion contains at least the first two sets of neurons involved in mechano-afferent signaling to the CNS.

Somatotopy in ascending pathways and other regions is found in a considerable diversity of species, both invertebrate^36–38^ and vertebrate^39–43^, and is a shared, fundamental organizing principle of nervous systems that permits efficient coding of incoming sensory information. The extraordinary complexity of the cephalopod body and its associated sensory systems has raised many questions about how large volumes of sensory information are encoded, but relatively few studies have successfully identified how such inputs are organized. Limited evidence for somatotopy has been found previously in various regions such as the fin and chromatophore lobes of the central brain through anatomical tracing studies^25,44^, whereas stimulation experiments of higher motor centers have demonstrated an apparent lack of somatotopy in centers associated with motor control an outputs^45,46^.

There are a number of interesting questions raised by the findings of primary afferent somatotopy here in this study. Due to the low-yield nature of the experiments (which could be improved significantly by future refinements), only gross mapping was demonstrated and individual neurons were not mapped relative to others within the same dermatome. Thus the precision and fine-scale organization of stellate ganglion somatotopy remains unclear at this stage. The temporal resolution of the imaging in this study was also very low; due to the curvature of the ganglion surface multiple imaging frames were needed to collect in-focus images of all cells involved in responses to touch. This limitation precluded temporal separation of the firing patterns of different cell classes, such as primary afferents and higher-order interneurons. Although retrograde labeling of cell bodies with axons projecting into the mantle nerves would likely have excluded higher-order neurons that have processes within the ganglion and not extending into the periphery, interneurons appear to be intermingled spatially with primary afferents, making either anatomical or functional separation challenging. More detailed anatomical mapping of the ganglion and molecular characterization of different neuronal sub-types is needed.

Currently, there are no genetically encoded calcium indicators (or any other genetic tools) in any cephalopod, and although the first attempts at gene editing have been successful^18^ in one squid species, the lack of tools available for live imaging is a considerable hinderance. Interest in cephalopods as comparative models of complex brains is growing rapidly along with new advances in culture and experimental methods, so the advent of genetically encoded markers of activity should greatly accelerate ongoing efforts to map functional properties of neural circuits in cephalopods. Rearing cephalopods through multiple generations, which is a necessity for establishing stable genetic lines, has always been exceptionally challenging. The emergence of the Euprymna genus (most notably *Euprymna berryi*) as a promising model for continuous culture, molecular studies and genetic manipulations is an important advance^47^. Although squid have a more restricted behavioral range than octopus, they have many advantages as model animals, including the large, accessible ganglia used in this study.

The mechanisms by which these mechanoreceptive neurons transduce touch are likewise unknown. Relatively few transcriptomic studies of the squid peripheral ganglia have been published. In a recent study of chemotactile receptors in the arm of octopuses, the mechanosensitive ion channel NompC, first described in Drosophila, was identified as being present in the sucker and arm tissue^48^. We are currently investigating whether this or other known mechanoreceptive ion channels are expressed in the dorsal-caudal region of the *Euprymna* stellate ganglion. Further molecular and transcriptomic characterization of these mechanoreceptive and mechano-nociceptive cells should help reveal further details of network properties and organization.

Cephalopods are known for their complex behavioral repertoires and advanced cognition^49,50^. Inherent in these abilities is the capacity for rapid and efficient processing of incoming sensory information. The role of the peripheral ganglia in the mantle and the arms of cephalopods is likely that of reflex responding, initial pre-processing, modulation and sorting of inputs such that only some information is transmitted to the central brain for whole-body behavioral responses. In this sense, the cephalopod peripheral ganglia are somewhat analogous to the vertebrate spinal cord. In vertebrates, somatotopy that begins in the spinal cord is preserved through ascending tracts and into cortical regions. Whether the same fundamental organizing principles of brain organization are present is cephalopods is a question of broad relevance for evolutionary and comparative neuroscience. This study provides the first functional evidence of somatotopy in cephalopods, and suggests that common principles of afferent neural circuit organization may have evolved independently in the complex, vertebrate-like nervous system of cephalopods.

## Acknowledgements and Funding

This study was conducted over the course of 5 years, including a period of maternity leave and the COVID-19 pandemic. I thank the various support systems that allowed ongoing, albeit slow, productivity on this extremely challenging project, most notably Dr. Ivan Anastassov - who provided invaluable practical help with microscopy, animal care and child care throughout. Funding was supplied by NSF IOS-2047331 and an Allen Distinguished Investigator award in Neural Circuit Design from the Frontiers Group of the Paul G. Allen Foundation. In-kind support was provided by the SFSU College of Science and Engineering in the form of teaching release and research-restart resources. Students and staff of the Crook lab from 2014-2022 provided animal care and husbandry.

## Figure legends

**Figure S1.**
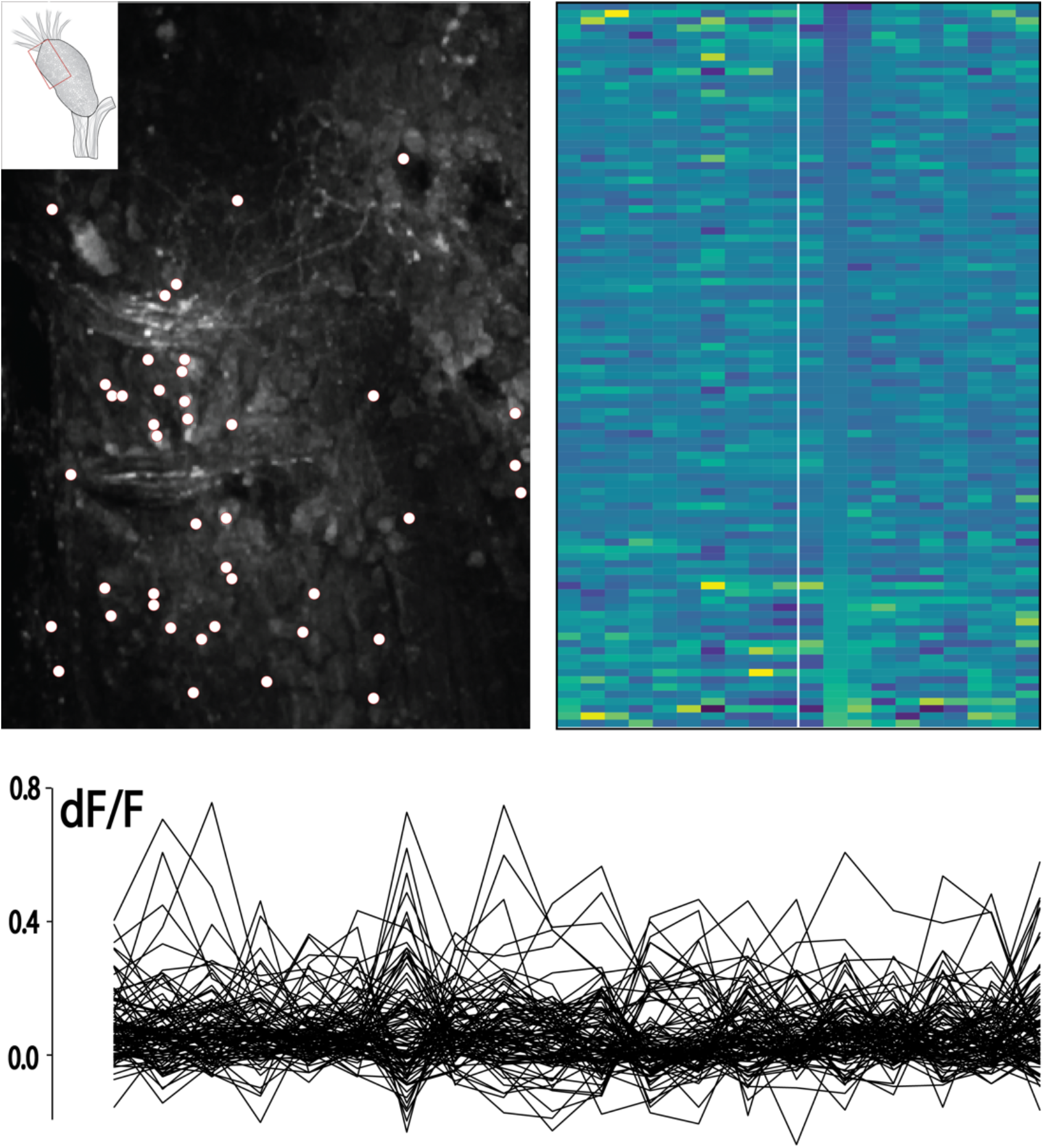
An example of analysis outputs from a ventral region on the ganglion, showing no clear responses to touch either in the heat-map plots or in the fluorescence traces from cells showing activity over the course of the imaging.

**Figure S2.**
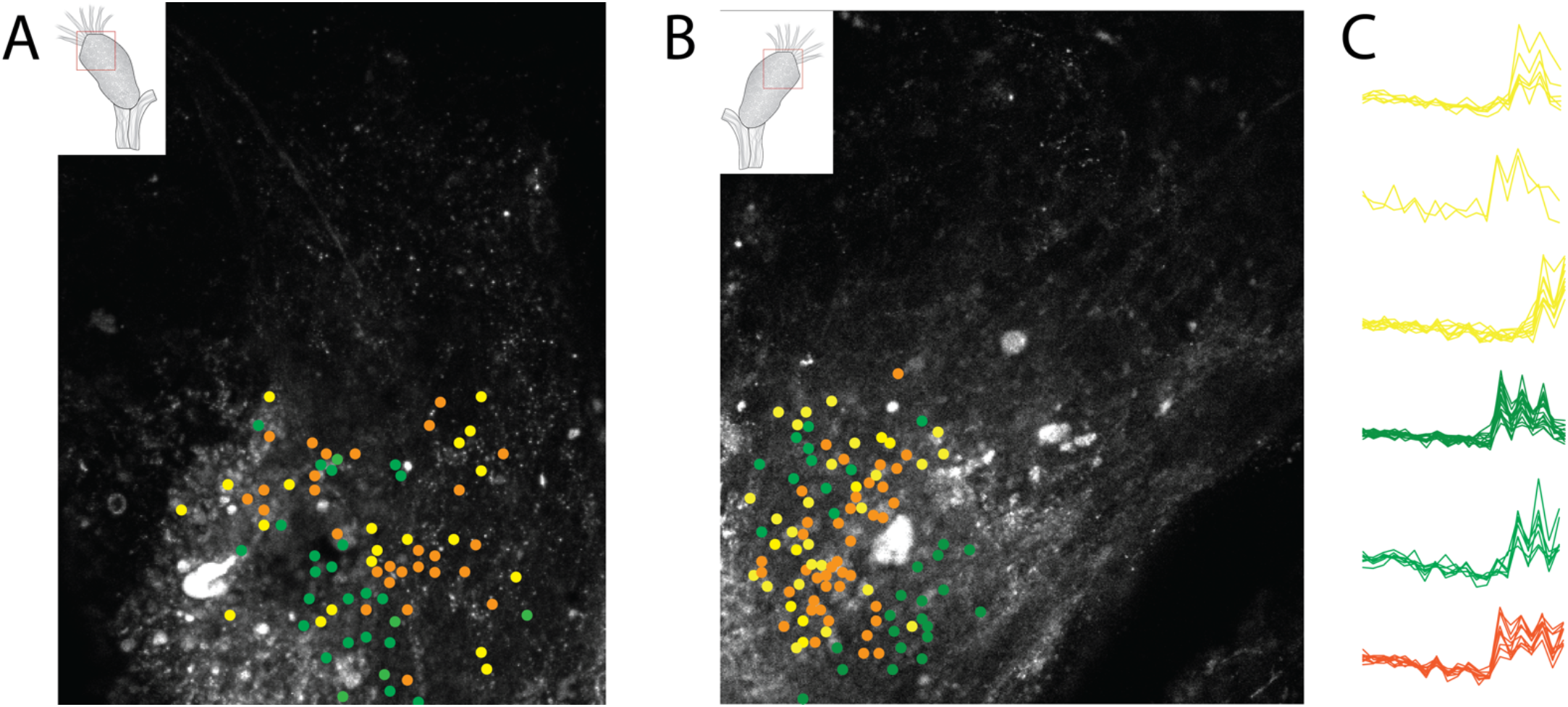
Interneuron-type cells plotted by tuning width suggests no clear organization.

Video S1. An example (corresponding to Figure panel 3A) of evoked activity recorded from the dorsal-caudal region of the stellate ganglion. Touches are applied to a different location on the ipsilateral mantle at frames 12, 14, 16, 18 and 20. Video has been stabilized and median-filtered in Fiji

Video S2. An example (corresponding to Figure panel 3B) of evoked activity recorded from the dorsal-caudal region of the stellate ganglion. Touches are applied to a different location on the ipsilateral mantle at frames 12, 14, 16, 18 and 20. Video has been stabilized and median-filtered in Fiji.

Video S3. An example recording from the ventral ganglion surface, showing no organized clustering of activity in response to mantle touch.

## Notes

### Competing Interest Statement

The authors have declared no competing interest.

